# EA-PheWAS: Integrating Phenotype Embeddings with PheWAS for Enhanced Gene-Phenotype Discovery

**DOI:** 10.64898/2026.04.21.720031

**Authors:** Wangjie Zheng, Tianyu Liu, Leqi Xu, Yuhan Xie, Yueqian Jing, Haoran Shao, Hongyu Zhao

**Affiliations:** Department of Biostatistics, Yale University, New Haven, CT, United States; Interdepartmental Program in Computational Biology and Bioinformatics, Yale University, New Haven, CT, United States

## Abstract

Phenome-wide association studies (PheWAS) enable systematic exploration of relationships between genetic variants and clinical phenotypes derived from electronic health records (EHRs). Conventional regression-based PheWAS treats phenotypes separately and relies on binary phenotype representations, which limits statistical power for rare variants and rare phenotypes and reduces the ability to detect associations with phenotypes that are distributed across clinical codes. To address this limitation, we first developed EmbedPheScan, a phenotype embedding–based prioritization framework that summarizes the phenotypic profiles of rare loss-of-function variant carriers in a continuous embedding space. We then proposed EA-PheWAS by combining these embedding-derived signals with conventional regression-based PheWAS results using the aggregated Cauchy association test. Using the UK Biobank whole-exome sequencing and EHR data, we show that the proposed methods maintain appropriate false-positive control. We then performed genome-wide phenome scans across all genes and across biologically defined gene classes to evaluate EA-PheWAS relative to conventional PheWAS and EmbedPheScan, consistently finding that EA-PheWAS outperformed the other two methods. We illustrate the utility of EA-PheWAS focusing on four genes representing distinct scenarios, including strong-effect disease genes (*PKD1, PKD2*), genes with large numbers of rare LoF carriers (*NF1*), and genes with extremely sparse carrier counts (*FBN1*).

## Introduction

Phenome-wide association studies (PheWAS) provide a systematic framework for identifying associations between genetic variants and a broad spectrum of clinical phenotypes derived from electronic health records (EHRs) [1]. Originally developed for common single-nucleotide polymorphisms (SNPs), PheWAS evaluates genotype–phenotype relationships by testing each phenotype separately using regression-based models, typically encoding disease phenotypes as binary indicators defined by diagnostic codes or phecodes. This paradigm enables genome-wide screening of associations and has been widely adopted in biobank-scale studies [2]. At the same time, the increasing availability of whole-exome sequencing (WES) data and longitudinal EHR data in biobanks has created new opportunities to investigate gene–phenotype relationships at much greater resolution.

PheWAS has been extended to rare variants through gene-level analyses, in which multiple rare variants within a gene are aggregated using burden [3] or kernel-based tests [4, 5] and evaluated against each phenotype [6]. This shift from common variants to rare variants is necessary because individual rare variants are observed in only a small number of individuals and therefore are typically underpowered for direct phenotype-wise association testing. In practice, rare-variant studies often focus on gene-level signals, particularly for rare loss-of-function (LoF) variants, which are more likely to have large phenotypic effects [7]. However, even in this setting, the overlap between rare variant carriers and phenotype cases is frequently too limited to support adequately powered association testing, even in large biobanks [6].

Despite these extensions, the PheWAS approach remains largely unchanged: phenotypes are treated as separate outcomes, and associations are assessed using covariate-adjusted regression models, typically logistic regression for binary traits. This formulation does not make use of the structure naturally encoded in longitudinal EHR which records capture phenotypes as sequences of ICD-10 diagnostic codes [8], and these codes exhibit hierarchical organization, semantic similarity, and strong co-occurrence patterns. However, conventional PheWAS collapses this rich structure into binary phenotype indicators, discarding information about similarity, hierarchy, and comorbidity. This limitation is particularly important for rare LoF carriers, whose clinical manifestations are often heterogeneous and pleiotropic, spanning multiple related phenotypes and organ systems [9]. In many cases, carriers of the same gene exhibit coherent phenotype patterns, but those signals are fragmented across multiple diagnostic codes, each of which may be too rare to achieve statistical significance when tested independently. As a result, regression-based PheWAS may fail to recover the broader and mechanistically coherent phenotype spectrum, instead prioritizing only the most prevalent or utilization-driven phenotypes [10].

Embedding models provide a principled framework for capturing latent relationships among clinical concepts by representing them as vectors, where proximity reflects similar clinical context. Methods such as Word2Vec [11] have been widely applied to EHR data to learn representations of medical codes, phenotypes, and patients [12–14]. These approaches have demonstrated strong performance in clustering prediction, and representation learning tasks, highlighting their ability to encode clinically meaningful structure. In our previous work, PERADIGM [14], we showed that embedding-similarity–based methods may substantially improve the identifications of disease-associated genes for rare diseases compared with conventional association methods using binary phenotypes such as burden tests [3], SKAT, and SKAT-O [4, 5]. These results suggest that embedding representations may capture clinically relevant relationships that are not accessible through conventional discrete phenotype representations.

Motivated by these observations, we extend the embedding-based method from rare-disease gene prioritization to PheWAS. We propose EA-PheWAS (Embedding-Augmented PheWAS), a unified framework that integrates signals from regression-based PheWAS and embedding-similarity–based phenotype prioritization. Rather than relying solely on phenotype-similarity-based ranking, EA-PheWAS leverages phenotype embeddings to complement conventional regression-based PheWAS, borrowing information from the embedding space to enhance the detection and prioritization of gene–phenotype associations. We first demonstrate through simulation studies that our method maintains appropriate control of the false positive rate. We then perform a genome-wide scan across all genes in the UK Biobank dataset to evaluate the overall performance of EA-PheWAS relative to conventional approaches. Next, we examine biologically informed gene subsets to evaluate the performance of EA-PheWAS in more functionally constrained gene sets rather than across all genes genome-wide. Finally, we report detailed case studies to characterize the conditions under which each method performs better and to illustrate how EA-PheWAS integrates complementary signals to improve phenotype prioritization. Collectively, these results demonstrate that EA-PheWAS provides a robust and effective framework for enhancing phenome-wide association analyses in large-scale population datasets.

## Results

### Overview of EA-PheWAS

As shown in Fig 1, EA-PheWAS integrates embedding-based phenotype similarity with regression-based PheWAS to enhance the discovery of gene–phenotype associations from EHR data and whole-exome sequencing data. The framework consists of three main components, which are described in detail in the Methods section.

**Fig 1.**
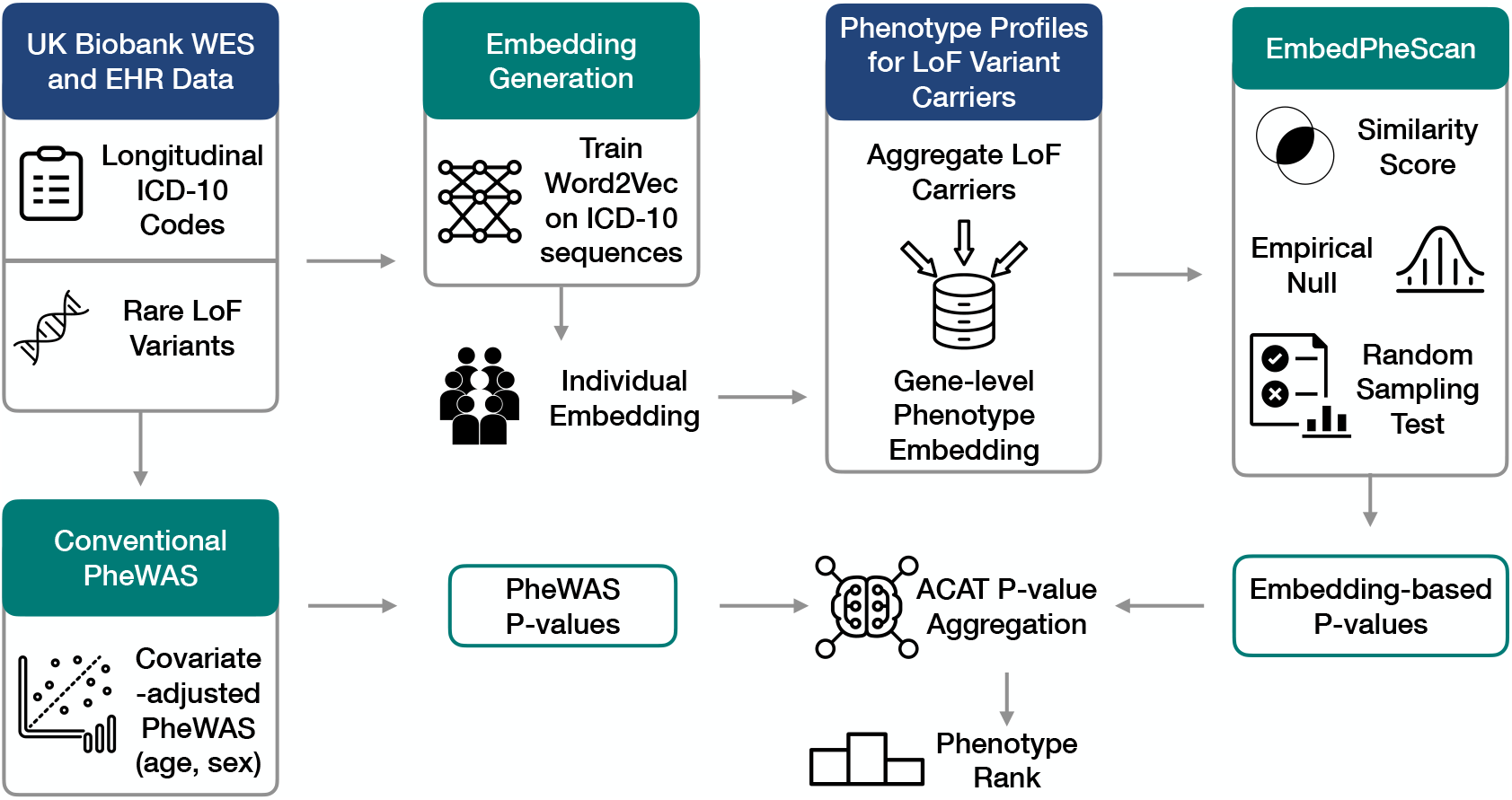
Workflow of EA-PheWAS. Longitudinal EHR data from the UK Biobank are used to train phenotype embeddings based on ICD-10 sequences. Individual-level phenotype embeddings are constructed and aggregated across rare loss-of-function (LoF) variant carriers to derive gene-level phenotype representations. In parallel, covariate-adjusted regression-based PheWAS is performed to obtain phenotype association p-values. Embedding-based similarity scores between gene-level embeddings and phenotype embeddings are converted into empirical p-values through null sampling. Finally, p-values from the embedding-based phenome scan and conventional PheWAS are combined using the ACAT method to produce aggregated association statistics, enabling improved prioritization of gene–phenotype associations.

First, phenotype embeddings are learned from longitudinal EHR data by training embedding models on sequences of ICD-10 codes. These embeddings capture latent relationships among phenotypes based on their co-occurrence patterns across individuals. The ICD-10 embeddings are mapped to phecodes, and individual-level phenotype embeddings are obtained by aggregating the embeddings of phecodes [1] observed in each individual’s medical records.

Second, we construct gene-level phenotype profiles by aggregating phenotype embeddings from individuals carrying rare LoF variants in each gene. This representation summarizes the phenotypic patterns observed among carriers and enables the quantification of gene–phenotype similarity. Based on this idea, we developed EmbedPheScan, an embedding-based phenome-wide phenotype prioritization framework that evaluates gene–phenotype relationships in the continuous phenotype embedding space. For each gene–phecode pair, a similarity is computed between the gene-level embedding and the phenotype embedding. An empirical null distribution is generated for each gene by repeatedly sampling random pseudo-carrier groups from the cohort, with each group matched to the observed number of rare LoF carriers for that gene. For each null replicate, we calculate the gene-level embedding from the sampled individuals and then calculate similarity scores between this null gene embedding and all phecode embeddings. Repeating this procedure 10,000 times produces an empirical null distribution for each gene–phecode pair, which is used to convert the observed similarity score into a standardized statistic and corresponding embedding-based p-value.

Third, EA-PheWAS integrates the complementary signals from embedding-based phenotype similarity and regression-based association testing. Specifically, p-values from the embedding-based phenome scan and the conventional PheWAS analysis are combined using the Aggregated Cauchy Association Test (ACAT). This aggregation step allows evidence from both approaches to contribute to the final association statistic, increasing sensitivity to phenotypes that may be weakly detectable by either method alone.

Compared with conventional regression-based PheWAS, EA-PheWAS leverages phenotype similarity captured by embeddings to borrow information across related clinical phenotypes. By combining embedding-derived evidence with conventional association testing, our proposed framework may improve the prioritization of gene–phenotype associations while retaining the statistical interpretability of p-value–based inference.

### EA-PheWAS can appropriately control false positive rates

To evaluate false-positive control, we performed simulations to compare EA-PheWAS, EmbedPheScan, and conventional regression-based PheWAS. We considered carrier counts of 50, 120, and 300 individuals in our simulations.

As shown in the QQ plots (Fig. 2), the three methods exhibited distinct behaviors when carrier counts are low. With 50 LoF carriers, conventional PheWAS produced a long tail of *p*-values equal to 1 because many phecodes had zero overlap between carriers and phenotype cases. With regression-based testing, this lack of overlap yielded non-informative association statistics. In contrast, EmbedPheScan does not rely solely on carrier–phenotype overlap and therefore reduces this degeneracy, although it showed mild inflation due to embedding-based similarity signals. By integrating both signals, EA-PheWAS balances these effects, producing a distribution that is less conservative than conventional PheWAS and less inflated than EmbedPheScan alone. When the carrier count increased to 120, all three approaches showed well-calibrated *p*-value distributions with similar tail behavior. In this case, both overlap-based association signals and embedding-based similarity could contribute to association tests. With 300 carriers, the proportion of p-values equal to 1 decreases substantially, reflecting the lower frequency of zero-overlap carrier–phenotype pairs under this null setting. Under this scenario, conventional PheWAS, EmbedPheScan, and EA-PheWAS all showed well-controlled null p-value distributions, indicating that none of the three methods generates excess false positives.

**Fig 2.**
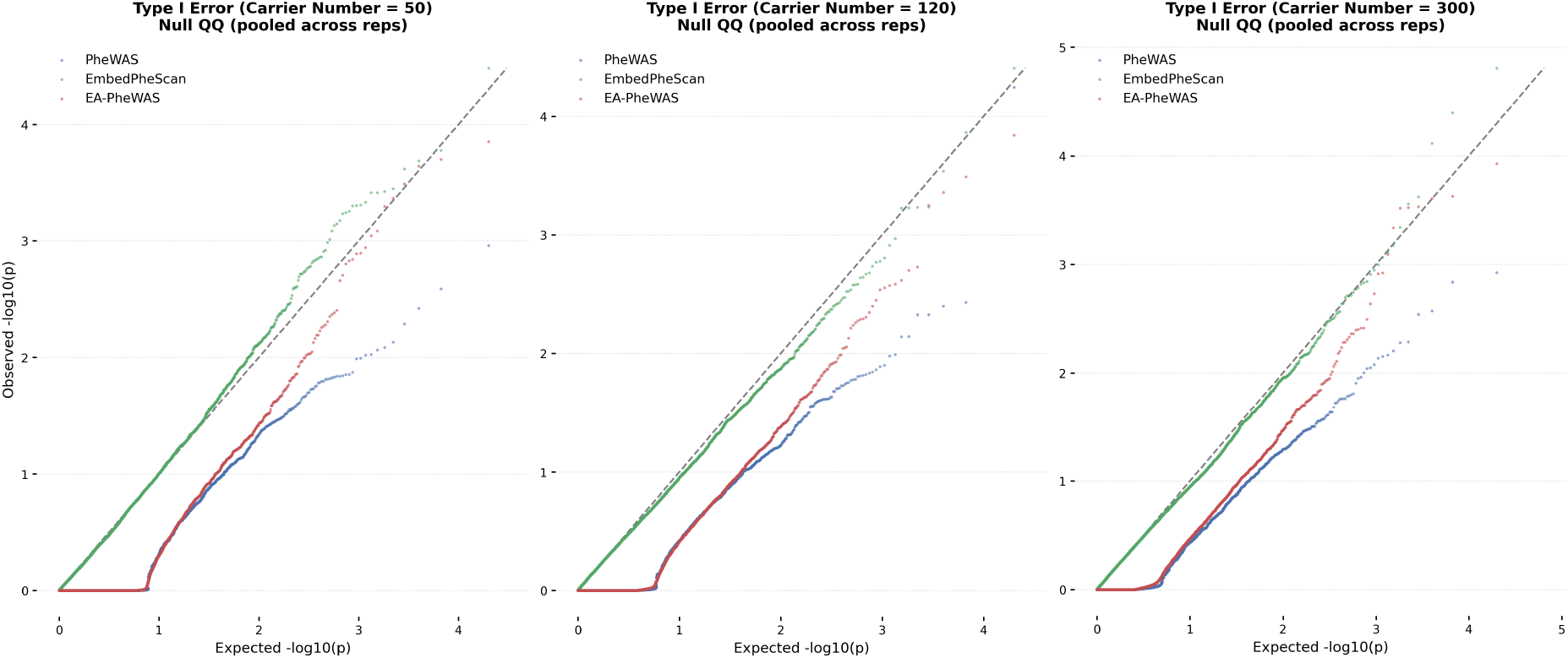
Null p-value distributions in type I error simulations. QQ plots of observed versus expected null *p*-values for conventional PheWAS, EmbedPheScan, and EA-PheWAS under simulated carrier counts of 50, 120, and 300. These carrier-count settings represent distinct rare-variant scenarios in the UK Biobank dataset.

We further summarize empirical type I error rates across multiple significance thresholds under different carrier-count settings. As shown in Fig. S1, all three methods maintained empirical false-positive rates close to the nominal levels across the null simulations. Together, these results indicate that EA-PheWAS preserves appropriate false-positive control across a range of carrier-count scenarios.

### EA-PheWAS can identify more trait-associated genes in the UKBB

To evaluate the genome-wide performance of EA-PheWAS, we applied it to all genes in the UK Biobank dataset and compared the results with conventional PheWAS and EmbedPheScan. Among the 17,233 genes tested, 4,545 genes have curated gene–phenotype relationships in the Human Phenotype Ontology (HPO) database and were therefore used as the reference set for evaluation (Fig. 3). These genes provide established phenotype annotations that enable quantitative comparison of phenotype prioritization performance across methods.

**Fig 3.**
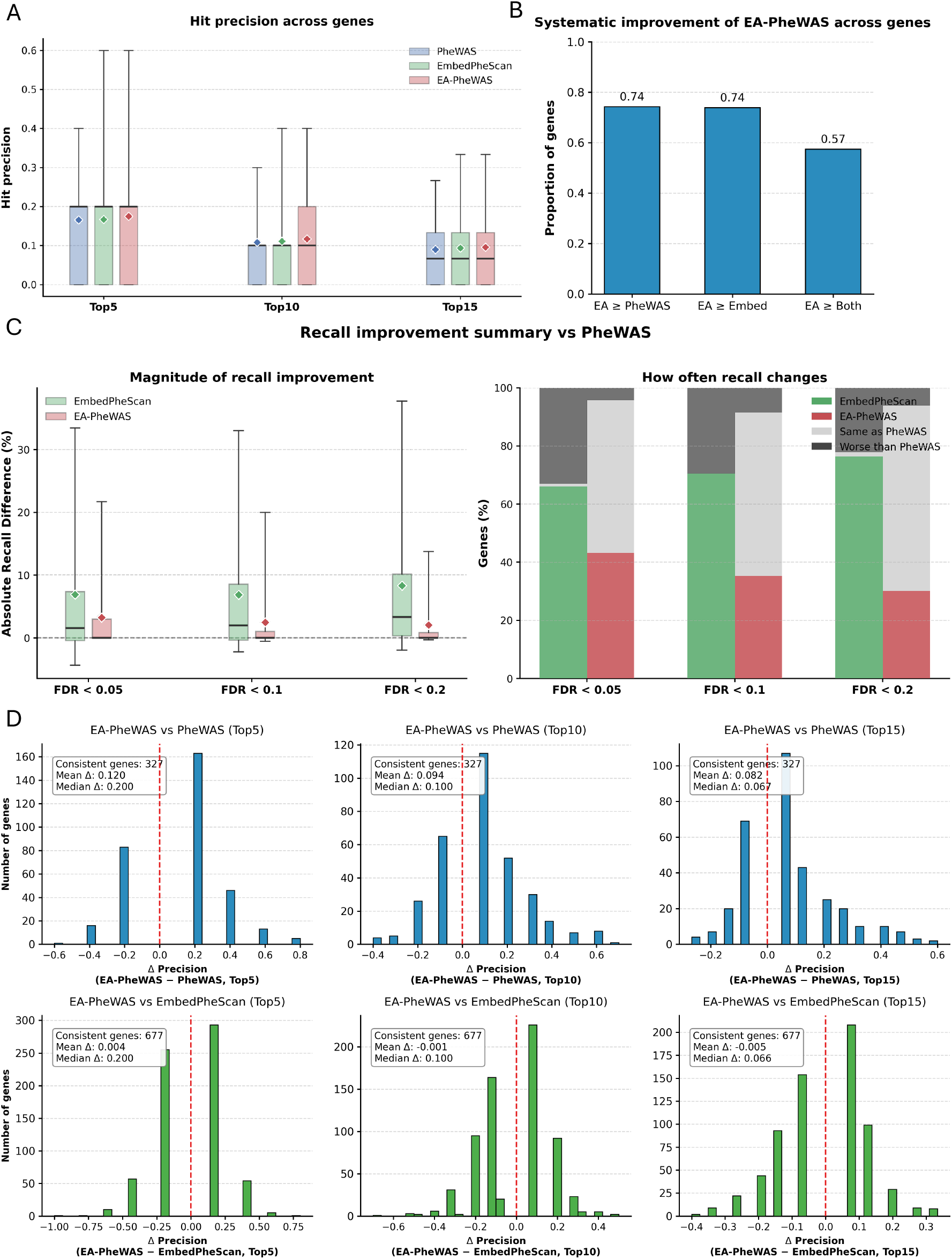
Genome-wide evaluation of phenotype prioritization performance in UK Biobank. (A) Hit precision across genes for conventional PheWAS, EmbedPheScan, and EA-PheWAS at Top-*k* phenotypes (*k* = 5, 10, 15). (B) Fraction of genes where EA-PheWAS achieves equal or higher hit precision than conventional PheWAS, EmbedPheScan, or both. (C) Absolute recall improvement relative to conventional PheWAS and the proportion of genes with improved recall across FDR thresholds. (D) Distribution of precision differences between EA-PheWAS and baseline methods for Top-*k* ranked phenotypes.

First, we evaluated hit precision across genes by comparing the average hit precision of the three methods at the Top-5, Top-10, and Top-15 ranked phenotypes (Fig. 3A and Table S1). Hit precision measures the proportion of top-*k* prioritized phenotypes that are also present in the HPO reference database. Across all three ranking thresholds, EA-PheWAS achieved the highest average hit precision, whereas EmbedPheScan consistently performed between EA-PheWAS and conventional PheWAS. Thus, both embedding-based approaches outperformed conventional PheWAS, with the integrated framework showing the greatest overall precision. These results indicate that EA-PheWAS can effectively combine complementary information from EmbedPheScan and conventional PheWAS to improve recovery of clinically relevant phenotypes among the top-ranked results.

Next, we evaluated recall, defined as the proportion of ground truth phenotypes recovered among the significant findings of each method. As shown in Fig. 3C and and Table S2, both EmbedPheScan and EA-PheWAS achieved higher recall than conventional PheWAS across multiple FDR thresholds, with EmbedPheScan showing the largest recall improvement. This is likely due to the fact that embedding-based similarity methods do not rely solely on direct carrier–phenotype overlap and can therefore identify related phenotypes even when the number of case carriers is small. By integrating embedding-based evidence with regression-based association testing, EA-PheWAS expands the set of detected phenotypes relative to conventional PheWAS while maintaining comparable overall behavior. To further examine this pattern, we quantified how often recall differed between methods across genes. As shown in Fig. 3C, across different FDR thresholds, EmbedPheScan and conventional PheWAS showed distinct recall patterns, indicating that the two methods capture complementary gene–phenotype information, while the recall differences between EA-PheWAS and conventional PheWAS are smaller, with many genes showing identical recall and a subset showing improved recall under EA-PheWAS. This pattern suggests that EA-PheWAS enhances phenotype discovery by borrowing information from EmbedPheScan while largely preserving the structure of conventional PheWAS results.

Finally, we examined the beat rate of EA-PheWAS relative to the other approaches. Rather than comparing average metrics alone, we identified genes for which one method consistently achieved higher hit precision across all Top-*k* settings (*k* = 5, 10, 15). As shown in Fig 3B, among genes with consistent ranking differences, EA-PheWAS outperformed conventional PheWAS, EmbedPheScan, and both methods simultaneously in most cases. To further characterize these improvements, we examined the distribution of precision differences for genes with consistent trends (Fig.3D), where Δprecision is defined as the difference in hit precision between EA-PheWAS and the comparison method at a given Top-*k* threshold. Across all Top-*k* settings, the distributions of Δprecision values are predominantly positive, indicating that EA-PheWAS improves phenotype ranking relative to both baseline methods for these genes.

Together, these genome-wide evaluations demonstrate that EA-PheWAS can effectively integrate embedding-based similarity with regression-based association testing. By borrowing complementary signals from EmbedPheScan while retaining the strengths of conventional PheWAS, EA-PheWAS may improve the prioritization of gene–phenotype associations in large-scale biobank analyses.

### EA-PheWAS performed well for haploinsufficient genes and tissue-specific genes

To further evaluate the ability of EA-PheWAS to identify gene–phenotype relationships, we next examined biologically informed gene subsets rather than all available genes. Specifically, we focused on two classes of genes with distinct functional properties: haploinsufficient (HI) genes and tissue-specific genes. Both groups are expected to exhibit more clinically coherent phenotype patterns and to be enriched for disease-relevant genes, providing a useful setting for testing whether EA-PheWAS is especially effective when gene effects are more functionally constrained and phenotypically concentrated.

We first investigated the HI gene group, which consists of genes for which loss of a single functional copy is sufficient to cause abnormal phenotypic consequences, and are therefore typically under strong functional constraint. We obtained a list of 417 HI genes from the ClinGen database [15]. Applying the same evaluation framework to this subset, we found that EA-PheWAS achieved the highest hit precision among the three methods, whereas EmbedPheScan remained intermediate between EA-PheWAS and conventional PheWAS (Fig.4A and Table S3). Notably, compared with the genome-wide evaluation across all genes, the separation between methods was larger in the HI subset, indicating that the advantage of EA-PheWAS becomes more pronounced in genes with more coherent and penetrant phenotypic effects. The recall analysis showed a similar pattern (Fig.4B): both EA-PheWAS and EmbedPheScan outperformed conventional PheWAS, while EA-PheWAS remained closer to conventional PheWAS overall but incorporated complementary findings from EmbedPheScan.

**Fig 4.**
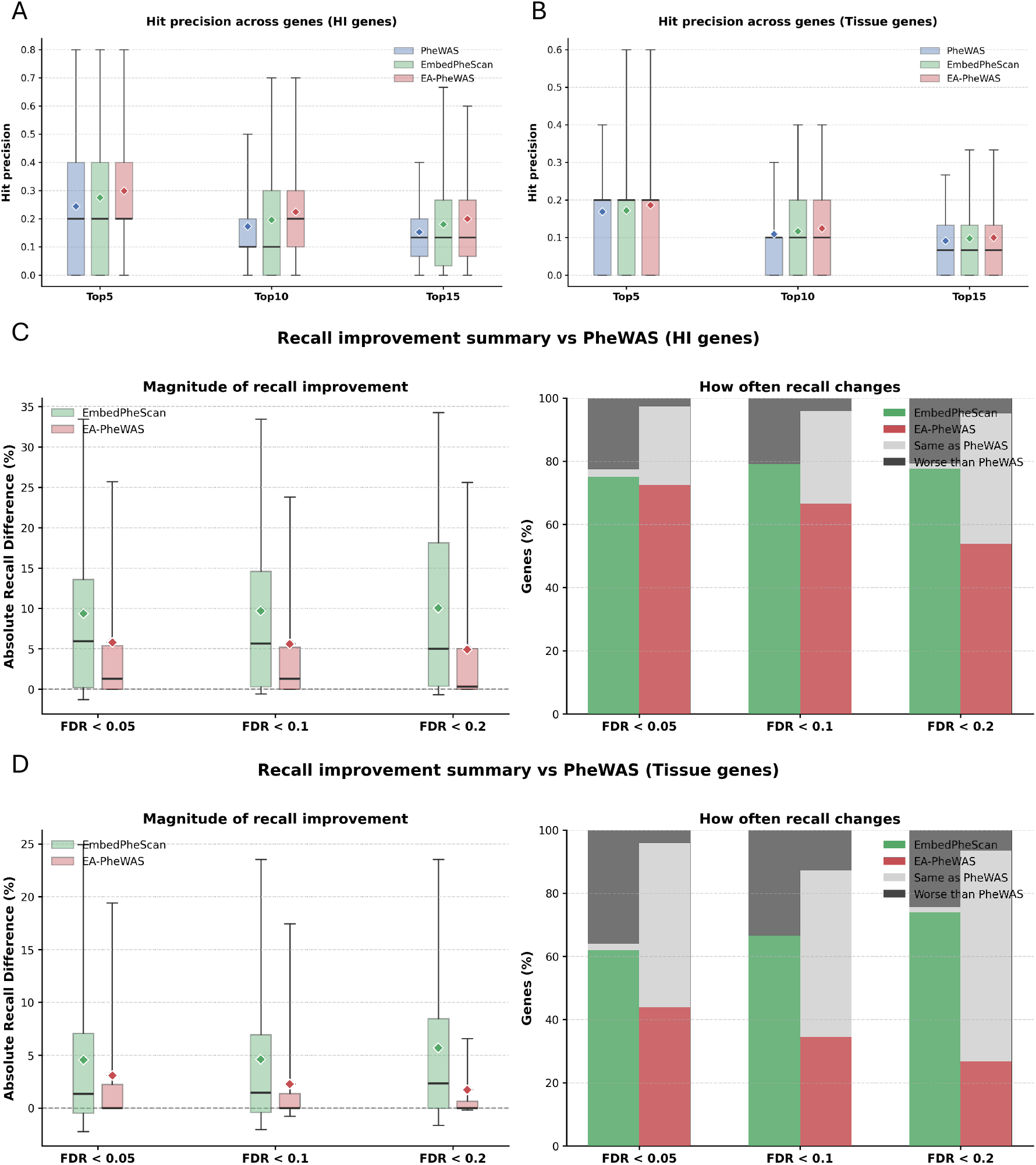
Genome-wide evaluation of phenotype prioritization performance across different functional gene subsets. (A) Hit precision across haploinsufficient (HI) genes for conventional PheWAS, EmbedPheScan, and EA-PheWAS at Top-5, Top-10, and Top-15 ranked phenotypes. (B) Hit precision across tissue-specific genes for conventional PheWAS, EmbedPheScan, and EA-PheWAS at Top-5, Top-10, and Top-15 ranked phenotypes. (C) Recall improvement summary relative to conventional PheWAS for HI genes, showing the magnitude of absolute recall differences and the proportion of genes with improved, unchanged, or worse recall across FDR thresholds. (D) Recall improvement summary relative to conventional PheWAS for tissue-specific genes, showing the magnitude of absolute recall differences and the proportion of genes with improved, unchanged, or worse recall across FDR thresholds.

We next investigated tissue-specific genes, which were obtained from the GTEx V8 resource [16]. Tissue-specific genes are genes whose expression is strongly enriched in particular tissues, and they are often associated with more localized biological functions and organ-specific disease processes. Because their effects are concentrated within specific tissues or systems, tissue-specific genes are expected to generate more structured phenotype patterns, making them particularly well suited for phenotype-similarity–based prioritization. Across 30 tissues, this set included 11,019 genes. Applying the same evaluation procedure to this subset again showed that EA-PheWAS achieved the highest hit precision among the three methods, with EmbedPheScan performing between EA-PheWAS and conventional PheWAS (Fig.4C and Table S4). Compared with tissue-specific genes, the improvement of EA-PheWAS was smaller in non–tissue-specific genes (Table S5), although it remained the best-performing method. Notably, EmbedPheScan underperformed conventional PheWAS in non–tissue-specific genes but outperformed it in tissue-specific genes, indicating that embedding-based prioritization is more effective for tissue-specific genes. The recall evaluation showed a similar trend (Fig.4D), with both EA-PheWAS and EmbedPheScan outperforming conventional PheWAS. These results indicate that embedding-based phenotype information remains informative for genes with tissue-restricted biological roles and that the integration step in EA-PheWAS can translate this structure into improved phenotype prioritization.

Together, these subgroup analyses strengthen the genome-wide findings by showing that EA-PheWAS performs particularly well in biologically constrained gene sets enriched for clinically coherent phenotypes. In both HI genes and tissue-specific genes, the improvement over the genome-wide baseline becomes more evident, supporting the idea that EA-PheWAS is especially effective when gene effects are more functionally specific and phenotype relationships are more structured. Beyond genome-wide evaluation, we performed gene-level case studies to examine how EA-PheWAS prioritizes phenotypes for individual genes and to better understand how the integration of embedding-based similarity and regression-based association testing improves gene-phenotype association discovery. We provide a few case studies in the following.

### Gene case study: *PKD1* and *PKD2*

We first focused on two major genes associated with autosomal dominant polycystic kidney disease (ADPKD), *PKD1* and *PKD2* [17–19]. ADPKD is characterized by progressive renal cyst formation and enlargement, ultimately leading to chronic kidney disease and end-stage renal failure. In addition to renal manifestations, ADPKD patients frequently exhibit hypertension, renal insufficiency, and complications involving the urinary tract and related organs [20, 21].

For *PKD1* (144 rare LoF variant carriers), the three methods had the same hit precision at the Top-5 level (Fig. 5B and Table S6) and EA-PheWAS achived the highest hit precision at the Top-10 and Top-15 levels. To investigate which phenotypes were uniquely identified by the aggregation step, we examined the detailed ranking comparison shown in Fig. 6A. The majority of the Top-15 phenotypes identified by EA-PheWAS are related to kidney and renal function, consistent with the known clinical manifestations of ADPKD. Notably, several phenotypes that are not explicitly listed in the HPO reference set, such as “kidney replaced by transplant” and “renal dialysis,” are nevertheless clinically associated with advanced ADPKD progression. The rank comparison further illustrates how EA-PheWAS integrates complementary signals from the two methods. For example, “End stage renal disease” is substantially uniquely identified relative to EmbedPheScan because conventional PheWAS detects strong carrier–phenotype overlap. Conversely, phenotypes such as “Vesicoureteral reflux” received substantially higher ranking relative to conventional PheWAS because embedding-based similarity captures clinically related phenotypes even when direct carrier–case overlap is limited. The differences among the Top-15 phenotypes across methods are summarized in Fig. 6B. EA-PheWAS not only uniquely identified phenotypes that are highly ranked by one method but overlooked by the other, but also identified phenotypes that are missed by both individual approaches. Among the phenotypes uniquely recovered by EA-PheWAS, “Acute glomerulonephritis” are present in the HPO ground truth set, indicating that the aggregation step can recover clinically validated phenotypes that are not strongly prioritized by either method alone. In contrast, although conventional PheWAS and EmbedPheScan individually identified additional phenotypes within the Top-15 list, a smaller proportion of these correspond to curated HPO annotations compared with EA-PheWAS. Finally, we examined the spatial distribution of prioritized phenotypes in the embedding space (Fig. 5A). Phenotypes identified by EmbedPheScan tend to cluster in the embedding space, reflecting the reliance of embedding-based methods on phenotype similarity. In contrast, conventional PheWAS identified phenotypes based primarily on carrier–phenotype overlap, which can produce more dispersed phenotypes. The phenotypes prioritized by EA-PheWAS exhibited an intermediate pattern, combining signals from both similarity-based clustering and overlap-based association testing.

**Fig 5.**
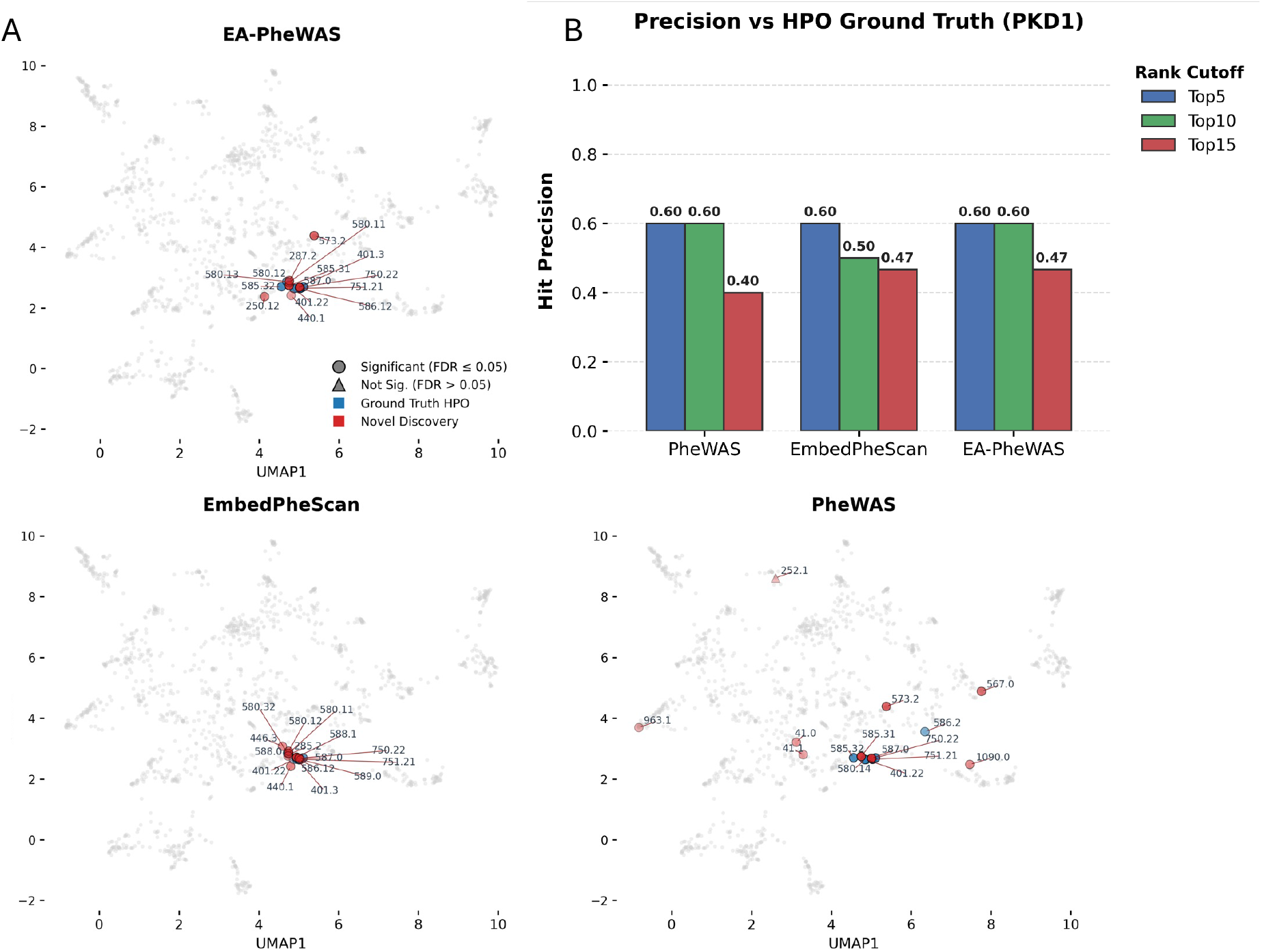
Case study for PKD1: embedding-space visualization and hit precision. (A) UMAP visualization of phenotype embeddings and prioritized phenotypes across methods. (B) Hit precision at different rank cutoffs.

**Fig 6.**
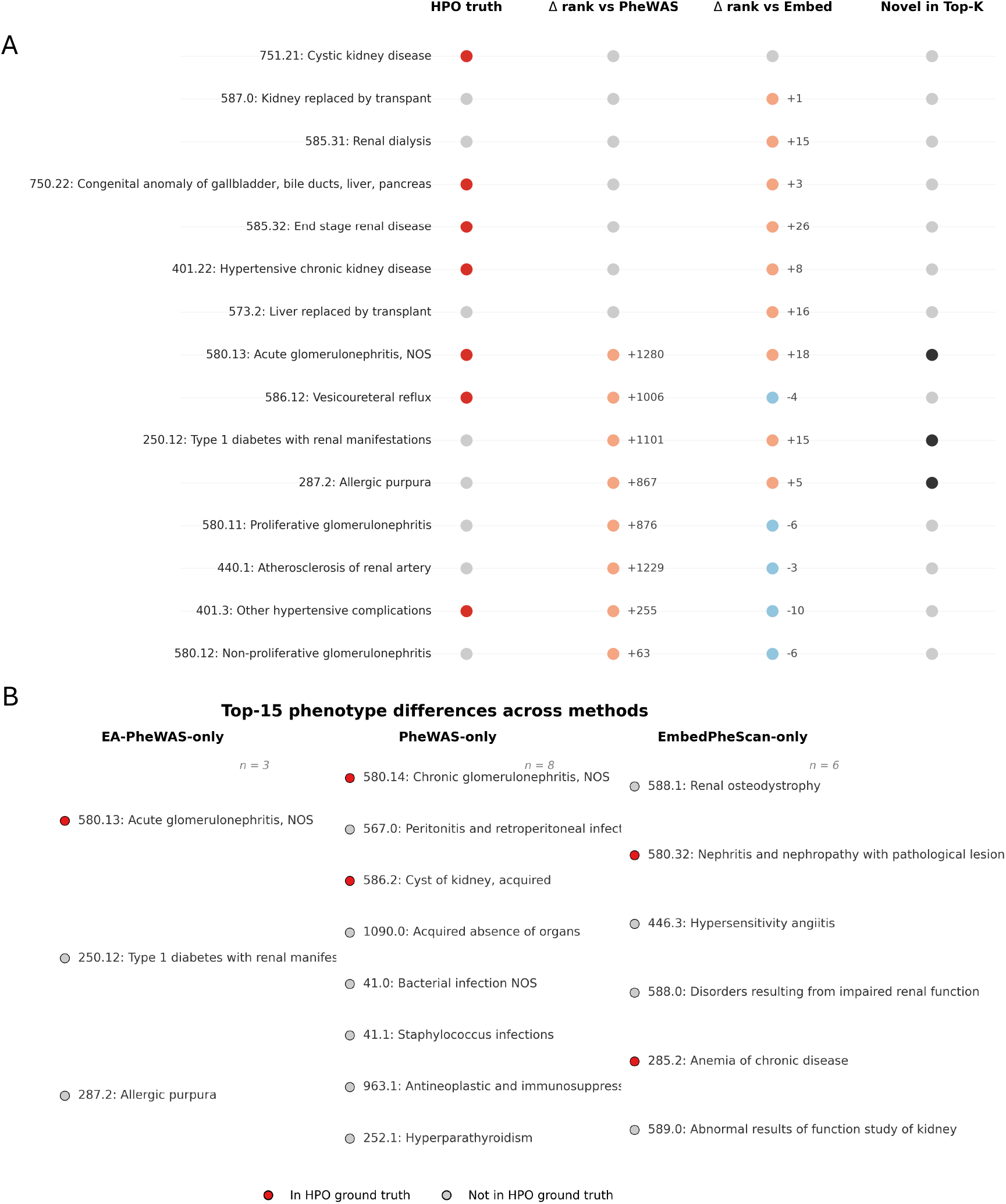
Case study for PKD1: phenotype ranking comparison across methods. (A) Comparison of top-ranked phenotypes across methods. Columns indicate whether the phenotype is present in the HPO ground truth, the change in ranking relative to conventional PheWAS (Δ rank vs PheWAS), the change in ranking relative to EmbedPheScan (Δ rank vs Embed), and whether the phenotype appears as a novel discovery within the top-*K* prioritized results. (B) Phenotypes appearing in the Top-15 results that differ across methods. Phenotypes are grouped into those uniquely identified by EA-PheWAS, those identified only by conventional PheWAS, and those identified only by EmbedPheScan. Markers indicate whether the phenotype is present in the HPO ground truth.

For *PKD2* (60 rare LoF variant carriers), EA-PheWAS consistently achieved higher hit precision than both conventional PheWAS and EmbedPheScan at the Top-5, Top-10, and Top-15 levels (Fig.7B and Table S7), indicating improved prioritization of *PKD2* -related phenotypes. Compared with the *PKD1* analysis, the advantage of EA-PheWAS was more pronounced for *PKD2*, likely reflecting the smaller number of *PKD2* carriers in the cohort and the resulting reduction in power for overlap-based association testing. The detailed ranking comparison (Fig.8A) further illustrates that EA-PheWAS uniquely identified phenotypes that are not strongly prioritized by either baseline method. Many of the phenotypes ranked near the top by EA-PheWAS originated from substantially lower rankings in both conventional PheWAS and EmbedPheScan. For example, phenotypes such as “Other hypertensive complications” and “Hypertensive chronic kidney disease” had limited direct overlap with *PKD2* carriers and were therefore ranked low by conventional PheWAS, but were substantially promoted by EA-PheWAS. This is biologically consistent with the well-established role of hypertension as a common manifestation of ADPKD [22]. In addition, several phenotypes not explicitly included in the HPO reference set for *PKD2*, including “kidney replaced by transplant” and “renal dialysis,” were highly prioritized by EA-PheWAS and are consistent with known complications of polycystic kidney disease. The Top-15 phenotype comparison across methods (Fig. 8B) showed that all the eight phenotypes uniquely uniquely identified by EA-PheWAS were present in the HPO reference set, whereas most phenotypes identified exclusively by conventional PheWAS or EmbedPheScan were not. Relative to the *PKD1* case study, the *PKD2* results provide stronger evidence that EA-PheWAS not only integrates signals from the two component methods, but can also recover clinically relevant phenotypes that are not prioritized by either method alone. Finally, examination of the embedding space (Fig. 7A) reveals distinct patterns for the two baseline methods. As in the *PKD1* example, phenotypes prioritized by EmbedPheScan form tight clusters in the embedding space, reflecting the reliance on phenotype similarity. In contrast, conventional PheWAS produces a more diffuse pattern, largely because the smaller number of *PKD2* carriers limits the statistical power of overlap-based testing, resulting in relatively few significant phenotypes after FDR correction. By integrating both sources of evidence, EA-PheWAS identified phenotypes both within the clustered similarity region and among surrounding phenotypes that are supported by association signals. Together, these results illustrate how EA-PheWAS improves phenotype prioritization for genes with weaker carrier signals by combining embedding-based similarity with regression-based evidence.

**Fig 7.**
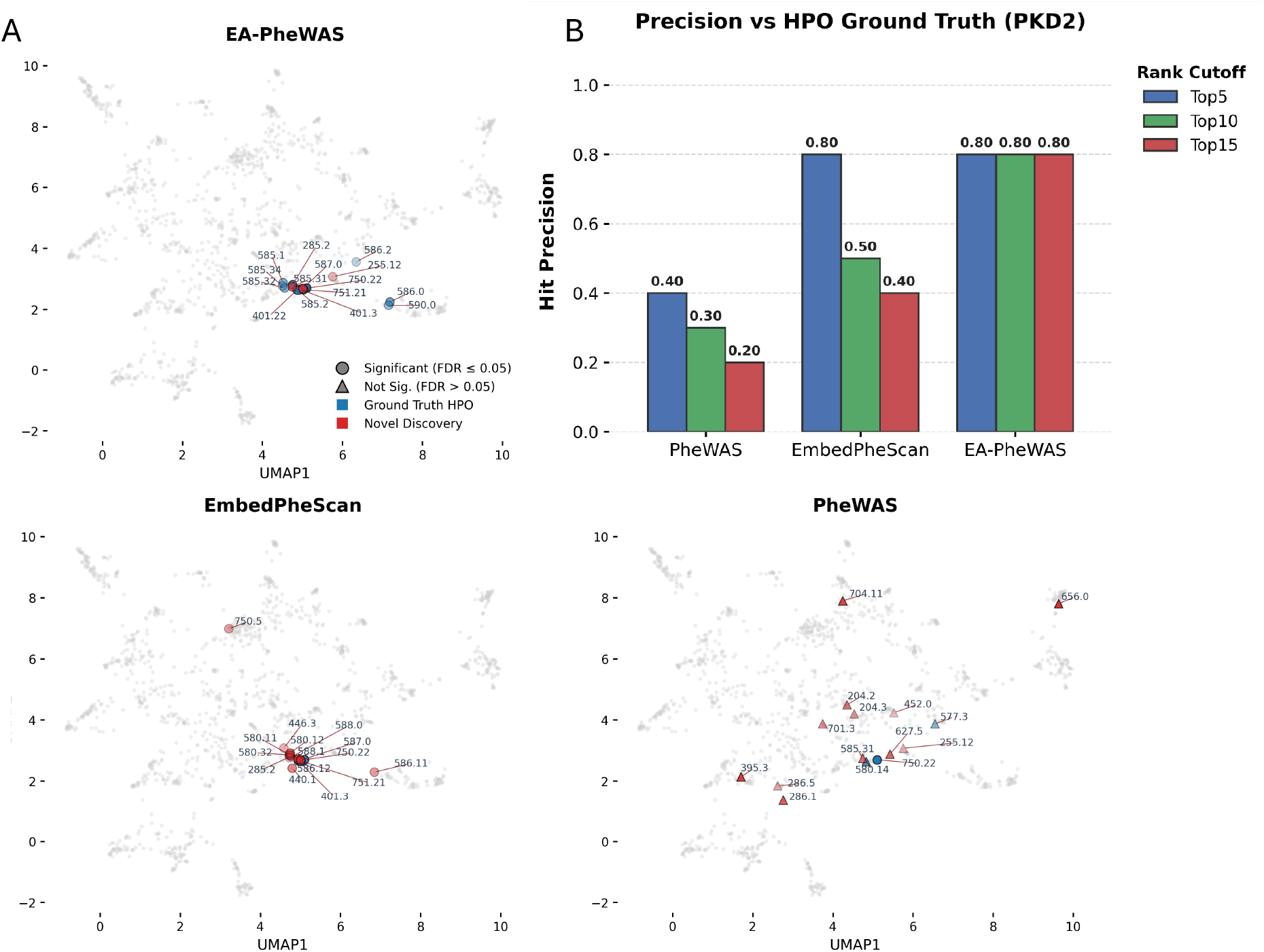
Case study for PKD2: embedding-space visualization and hit precision. (A) UMAP visualization of phenotype embeddings and prioritized phenotypes across methods. (B) Hit precision at different rank cutoffs.

**Fig 8.**
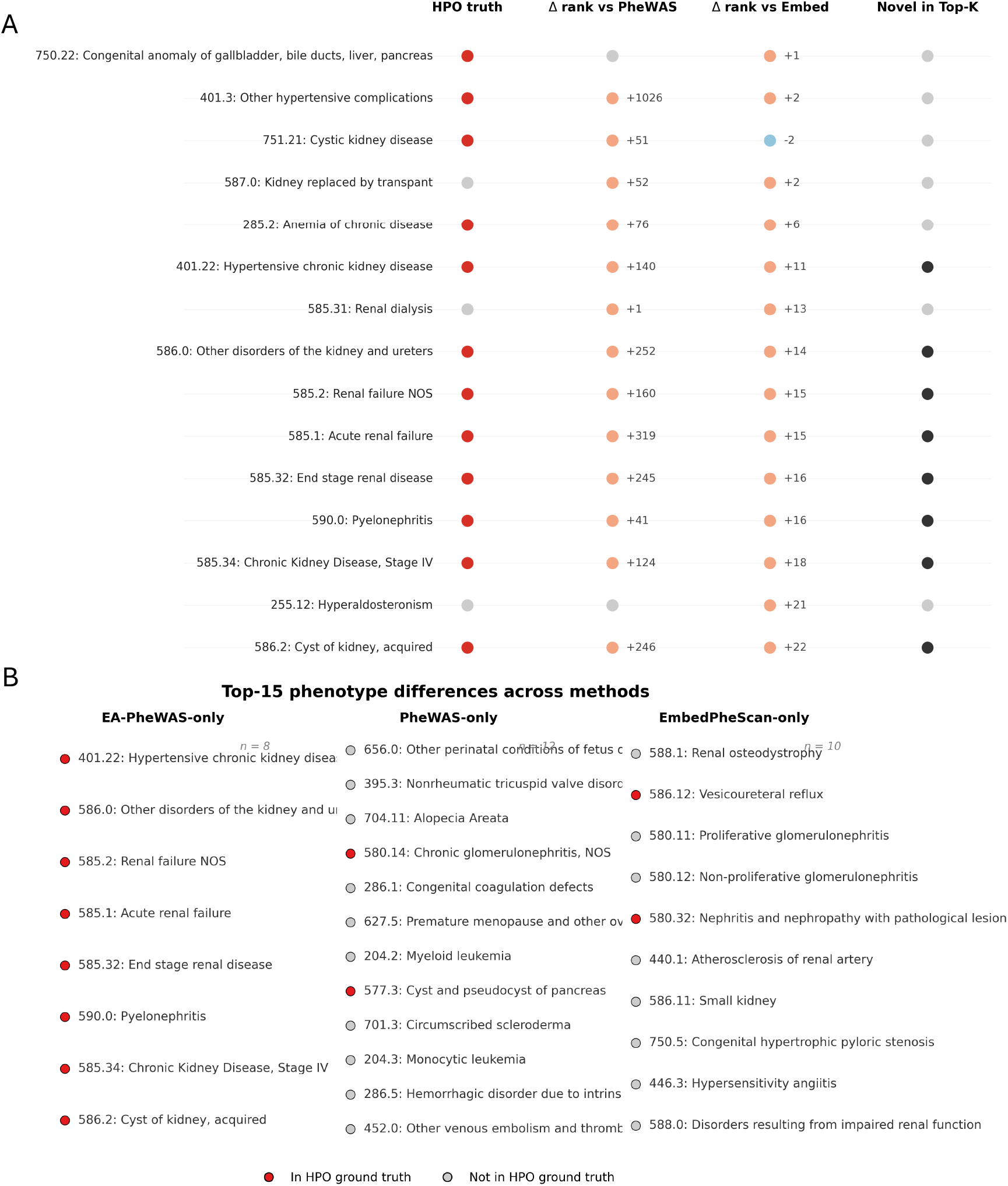
Case study for PKD2: phenotype ranking comparison across methods. (A) Comparison of top-ranked phenotypes across methods. Columns indicate whether the phenotype is present in the HPO ground truth, the change in ranking relative to conventional PheWAS (Δ rank vs PheWAS), the change in ranking relative to EmbedPheScan (Δ rank vs Embed), and whether the phenotype appears as a novel discovery within the top-*K* prioritized results. (B) Phenotypes appearing in the Top-15 results that differ across methods. Phenotypes are grouped into those uniquely identified by EA-PheWAS, those identified only by conventional PheWAS, and those identified only by EmbedPheScan. Markers indicate whether the phenotype is present in the HPO ground truth.

### Gene case study: *NF1*

Next, we examined *NF1*, which has a relatively large number of rare loss-of-function (LoF) carriers in our cohort (311 carriers). *NF1* encodes neurofibromin, a tumor suppressor that negatively regulates RAS signaling, and pathogenic variants in *NF1* are associated with neurofibromatosis type 1, a multisystem disorder characterized by cutaneous, neurological, and neoplastic manifestations [23–25]. Because of its relatively large carrier count and broad clinical spectrum, *NF1* provides an informative setting for evaluating the performance of EA-PheWAS in a gene with substantial overlap-based signal as well as complex phenotype structure.

As shown in Fig. 9B and Table S8, EA-PheWAS achieved the highest hit precision across all Top-*K* settings, while conventional PheWAS performed relatively well at Top-5 and Top-10 and EmbedPheScan showed stronger performance at Top-10 and Top-15. This pattern suggests that, with sufficient carrier counts, conventional PheWAS can effectively prioritize phenotypes supported by direct carrier–phenotype overlap, whereas EmbedPheScan captures additional phenotypes through similarity in the embedding space. By integrating these two complementary signals, EA-PheWAS achieved the most accurate overall ranking. The detailed comparison of top-ranked phenotypes (Fig. 10A) further illustrates this integrative behavior. The highest-ranked phenotypes under EA-PheWAS were largely driven by strong signals from conventional PheWAS, whereas several additional top phenotypes appeared to be promoted primarily through EmbedPheScan. For example, “benign neoplasm of brain, cranial nerves, meninges” was not strongly prioritized by conventional PheWAS because of limited direct overlap with *NF1* carriers, but was highly ranked by EmbedPheScan because of its proximity to other neuro-related phenotypes in the embedding space. More broadly, multiple phenotypes exhibited substantial rank shifts relative to one or both baseline methods, indicating that EA-PheWAS effectively integrates overlap-based association evidence with phenotype-similarity information. Consistent with this pattern, Fig. 10B shows that EA-PheWAS uniquely identified five phenotypes, three of which are present in the HPO reference set. These include phenotypes such as “cancer of connective tissue,” “facial nerve disorders,” and “hydrocephalus,” all of which are biologically plausible in the context of *NF1* -related neurological and tissue manifestations [26–28]. In contrast, although conventional PheWAS and EmbedPheScan also identified method-specific phenotypes, the proportion of HPO-supported phenotypes among these unique findings was lower than for EA-PheWAS. The embedding-space visualization further illustrates the complementary nature of the three methods (Fig. 9A). Compared with the PKD1 and PKD2 case studies, the prioritized phenotypes for *NF1* were more dispersed, consistent with the broader involvement of both neurological and tissue-related systems. Phenotypes identified by conventional PheWAS were widely separated in the embedding space, whereas those prioritized by EmbedPheScan were concentrated in two main clusters. EA-PheWAS captured features of both patterns, generating a phenotype profile that reflects both embedding-based similarity and overlap-based association. Together, these results show that for genes with relatively large carrier counts and complex multisystem phenotypes, EA-PheWAS can effectively combine the strengths of conventional PheWAS and EmbedPheScan to improve phenotype prioritization

**Fig 9.**
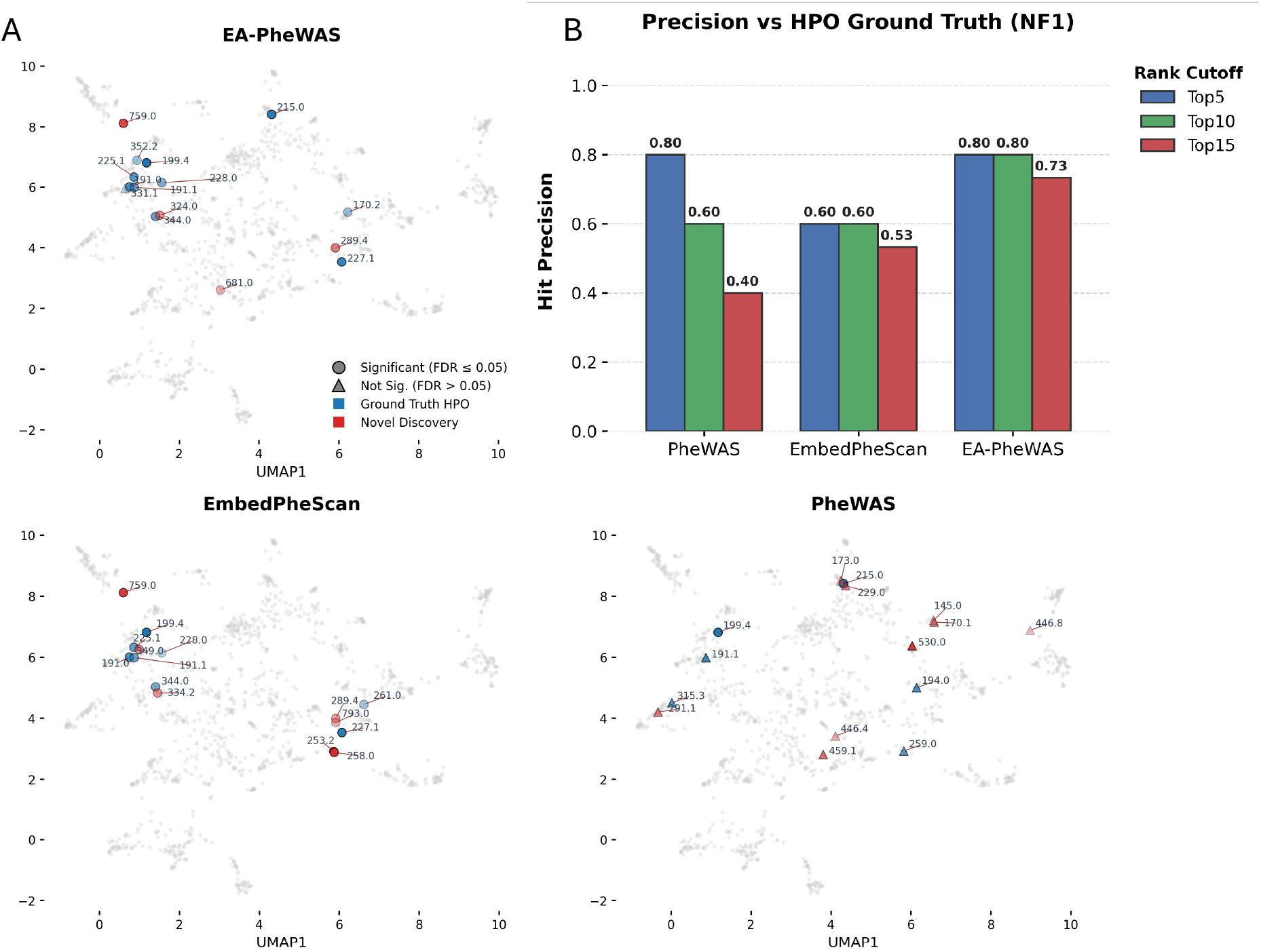
Case study for NF1: embedding-space visualization and hit precision. (A) UMAP visualization of phenotype embeddings and prioritized phenotypes across methods. (B) Hit precision at different rank cutoffs.

**Fig 10.**
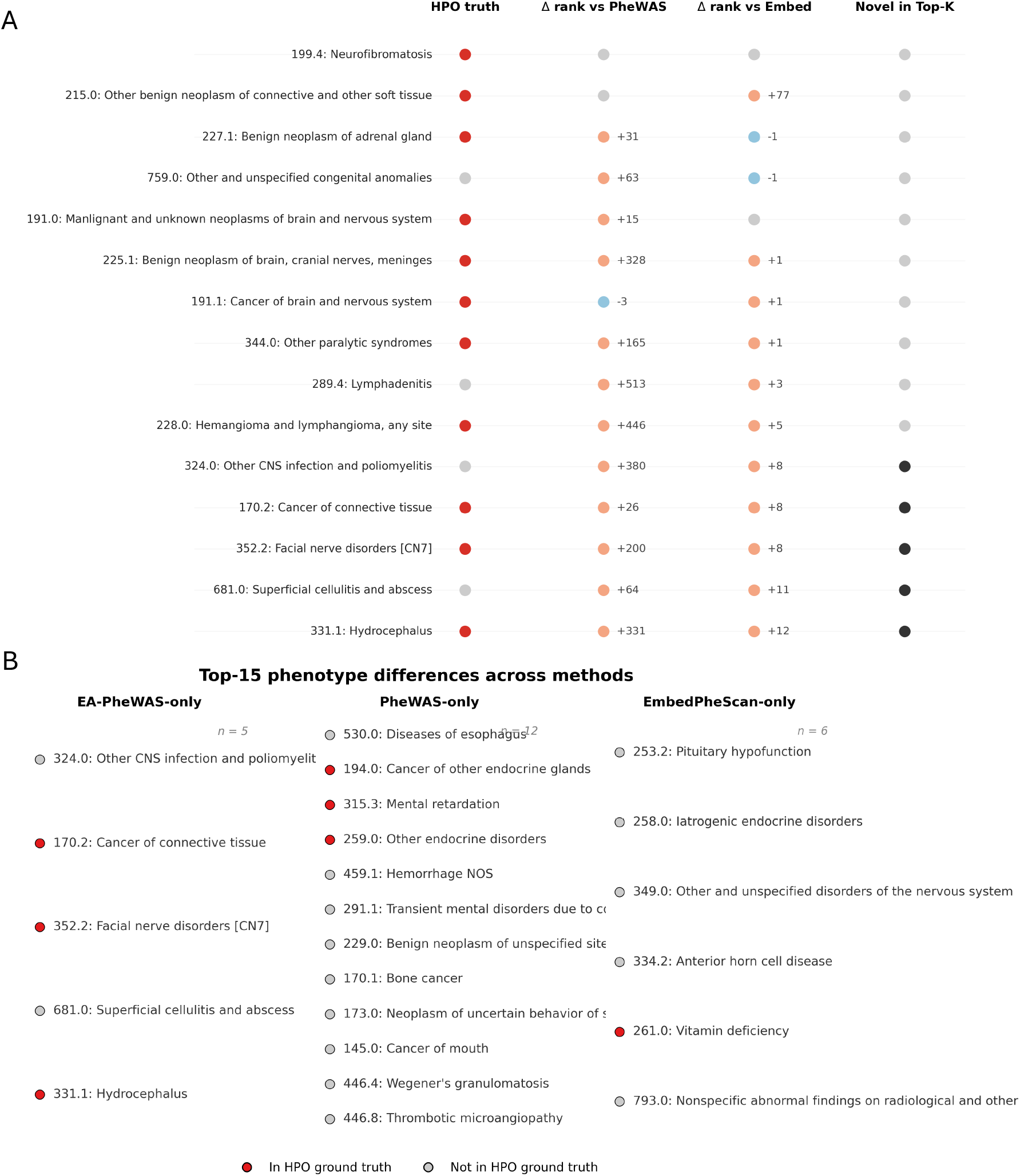
Case study for NF1: phenotype ranking comparison across methods. (A) Comparison of top-ranked phenotypes across methods. Columns indicate whether the phenotype is present in the HPO ground truth, the change in ranking relative to conventional PheWAS (Δ rank vs PheWAS), the change in ranking relative to EmbedPheScan (Δ rank vs Embed), and whether the phenotype appears as a novel discovery within the top-*K* prioritized results. (B) Phenotypes appearing in the Top-15 results that differ across methods. Phenotypes are grouped into those uniquely identified by EA-PheWAS, those identified only by conventional PheWAS, and those identified only by EmbedPheScan. Markers indicate whether the phenotype is present in the HPO ground truth.

### Gene case study: *FBN1*

Finally, we examined *FBN1*, which has only 18 rare loss-of-function (LoF) carriers in our dataset and therefore provides a stringent example for evaluating EA-PheWAS under extremely sparse carrier conditions. *FBN1* encodes fibrillin-1, a major structural component of extracellular microfibrils that is critical for connective tissue integrity and elastic fiber assembly. Pathogenic variants in *FBN1* are the primary cause of Marfan syndrome, a multisystem disorder affecting the cardiovascular, skeletal, and ocular systems, with well-recognized manifestations including aortic disease and valvular abnormalities [29, 30].

As shown in Fig. 11B and Table S9, EA-PheWAS achieved the highest hit precision across all Top-*k* settings. In contrast to genes with larger carrier counts, conventional PheWAS performed relatively poorly for *FBN1*, whereas EmbedPheScan showed stronger phenotype prioritization. This pattern is consistent with the extremely small number of carriers, under which regression-based association testing has limited power because of the lack of direct carrier–phenotype overlap. As a result, EA-PheWAS relied more heavily on the signal contributed by EmbedPheScan. The detailed ranking comparison (Fig. 12A) supports this interpretation. Overall, the phenotype ranking produced by EA-PheWAS more closely resembled that of EmbedPheScan than that of conventional PheWAS, with the exception of several very strong signals such as “Chromosomal anomalies and genetic disorders” and “Diseases of tricuspid valve,” which were also strongly supported by conventional PheWAS. Several biologically relevant phenotypes with limited overlap in carriers were not highly prioritized by conventional PheWAS but were promoted by EA-PheWAS through the embedding-based signal. For example, “Mitral valve disease” had limited direct overlap with *FBN1* LoF carriers and was therefore difficult to detect by conventional PheWAS, but it shares high similarity with other cardiovascular and valvular phenotypes in the embedding space and has strong biological support in the context of *FBN1* -related disease [31]. These results indicate that, for *FBN1*, EA-PheWAS benefits primarily from the phenotype-similarity information captured by EmbedPheScan. Importantly, EA-PheWAS did not simply reproduce the results of either baseline method. As shown in Fig. 12B, the three phenotypes uniquely identified by EA-PheWAS were all present in the HPO reference set, whereas the proportion of HPO-supported phenotypes among method-specific findings was lower for both conventional PheWAS and EmbedPheScan. Notably, compared with conventional PheWAS, EmbedPheScan recovered a higher proportion of true phenotypes among its unique findings, further supporting the value of embedding-based prioritization when carrier numbers are extremely limited. The embedding-space visualization further illustrates these differences (Fig. 11A). Phenotypes identified by EA-PheWAS and EmbedPheScan formed relatively compact and coherent clusters, whereas those identified by conventional PheWAS were more dispersed in the embedding space. Together, these results show that when carrier counts are extremely low, EA-PheWAS continues to integrate both sources of evidence but derives greater benefit from the embedding-based component, enabling recovery of clinically relevant phenotypes that are difficult to detect through overlap-based association testing alone.

**Fig 11.**
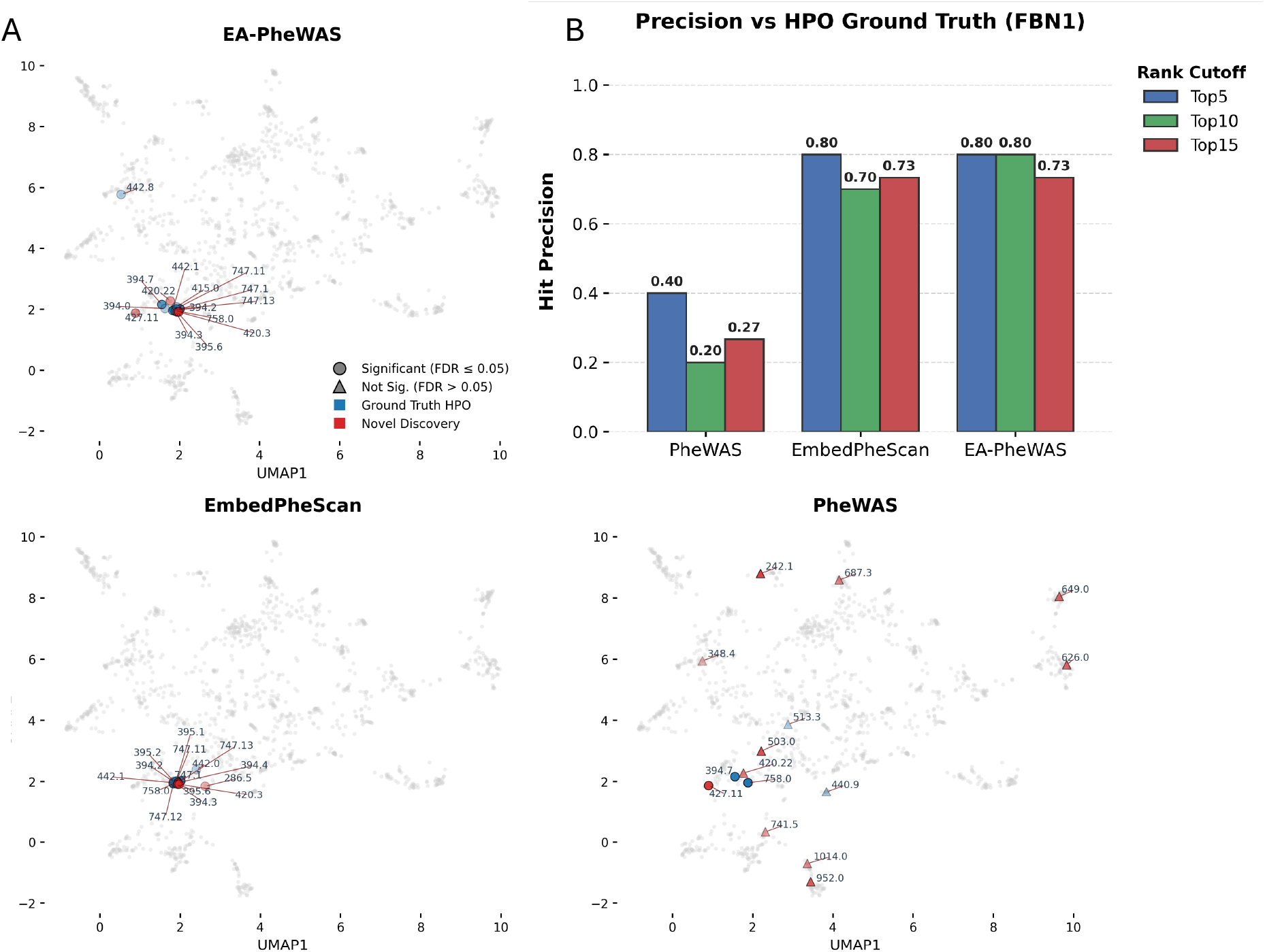
Case study for FBN1: embedding-space visualization and hit precision. (A) UMAP visualization of phenotype embeddings and prioritized phenotypes across methods. (B) Hit precision at different rank cutoffs.

**Fig 12.**
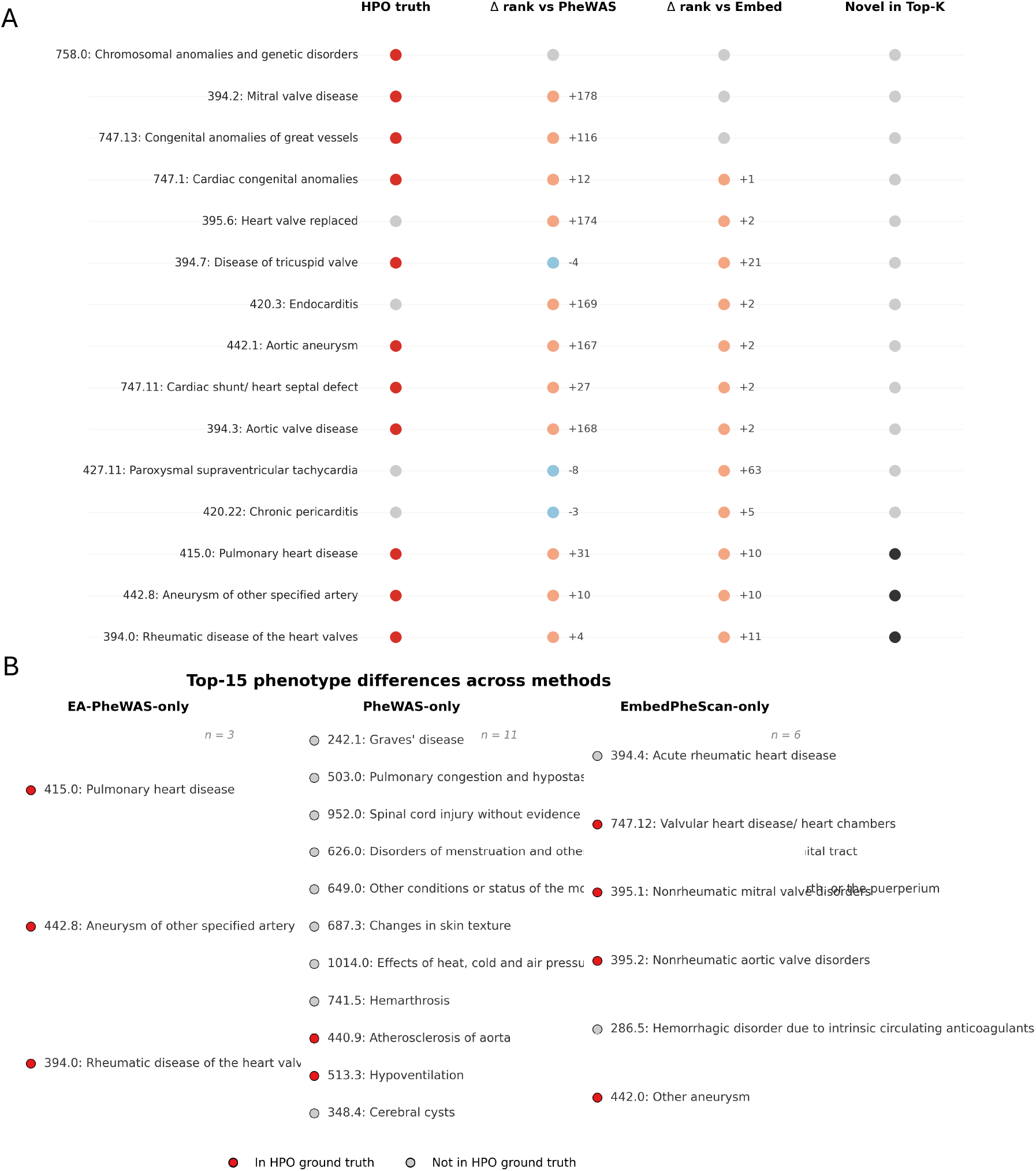
Case study for FBN1: phenotype ranking comparison across methods. (A) Comparison of top-ranked phenotypes across methods. Columns indicate whether the phenotype is present in the HPO ground truth, the change in ranking relative to conventional PheWAS (Δ rank vs PheWAS), the change in ranking relative to EmbedPheScan (Δ rank vs Embed), and whether the phenotype appears as a novel discovery within the top-*K* prioritized results. (B) Phenotypes appearing in the Top-15 results that differ across methods. Phenotypes are grouped into those uniquely identified by EA-PheWAS, those identified only by conventional PheWAS, and those identified only by EmbedPheScan. Markers indicate whether the phenotype is present in the HPO ground truth.

## Discussion

Investigating phenotypes associated with a given gene has long been a central goal of human genetics, and PheWAS has become a key framework for systematically characterizing gene–phenotype relationships. As whole-exome and whole-genome sequencing data become increasingly available, an important challenge is how to make effective use of rare variant information for phenome-wide analyses. Unlike conventional SNP-based PheWAS, which is typically well powered because common variants are observed in a large number of individuals, rare-variant PheWAS generally requires gene- or region-level aggregation to combine multiple variants into a burden signal. However, the underlying statistical framework remains largely unchanged: phenotypes are still represented as binary traits and tested independently, with little consideration of the relationships among phenotypes. At the same time, the growing availability of HER data provides rich individual-level phenotype information that captures longitudinal and correlated clinical patterns. Motivated by the observation that rare variant carriers often exhibit coherent phenotype patterns, we developed EA-PheWAS, an embedding-augmented PheWAS framework that integrates phenotype similarity information with conventional association testing to improve gene–phenotype discovery.

In EA-PheWAS, we combine two complementary sources of evidence using ACAT. The first is the conventional regression-based PheWAS signal, which primarily captures associations through direct overlap between phenotype cases and rare LoF carriers of a given gene and performs well when carrier counts and phenotype prevalence are sufficiently large. The second is the embedding-similarity signal from EmbedPheScan, which prioritizes phenotypes according to similarity in the learned phenotype embedding space and therefore does not rely solely on direct carrier–phenotype overlap. Instead, it leverages semantic and clinical relatedness among phenotypes, allowing recovery of phenotypes that may be weakly represented in sparse carrier groups. By integrating these two signals, EA-PheWAS can improve phenotype prioritization for both relatively common and rare phenotypes associated with a given gene.

This complementary behavior was evident in both the genome-wide evaluation and the gene-level case studies. EA-PheWAS consistently improved hit precision and recall relative to both conventional PheWAS and EmbedPheScan across the full gene set as well as across distinct functional gene subsets. For *PKD1* and *PKD2*, which represent strong-effect genes for a specific disease, EA-PheWAS recovered phenotypes related to glomerulonephritis, hypertension, and disease progression by combining direct association evidence with clinically related phenotypes in the embedding space. For *NF1*, which has a relatively large number of rare LoF carriers, EA-PheWAS benefited substantially from the strong overlap-based signal detected by conventional PheWAS while incorporating additional biologically plausible phenotypes from EmbedPheScan. In contrast, for *FBN1*, where the number of carriers was extremely limited, EA-PheWAS relied more heavily on embedding-based similarity and was able to recover heart relevant phenotypes that were weakly supported by regression alone. Across these settings, EA-PheWAS not only integrated phenotypes prioritized by either method individually, but also recovered phenotypes that were not strongly highlighted by either approach alone, including phenotypes supported by HPO-based ground truth.

Although EA-PheWAS shows strong performance in rare-variant PheWAS using WES data, several areas remain for further development. First, our analyses were performed in individuals of European ancestry to minimize potential confounding from population structure and to provide a well-controlled initial evaluation. Extending the framework to multi-ancestry settings will be an important next step for improving generalizability. Second, we used Word2Vec-based phenotype embeddings to represent individual phenotype profiles. This approach performed well in our analyses and provided an effective starting point, although future work may explore alternative embedding architectures that capture additional aspects of longitudinal clinical structure. Third, the current study focuses on the UK Biobank, which offers a large and well-characterized resource for method development and evaluation. Applying EA-PheWAS to additional biobank-scale datasets and external cohorts will further clarify its robustness and portability across populations and healthcare systems.

In summary, EA-PheWAS provides a complementary perspective for phenome-wide association analysis by integrating regression-based association testing with embedding-based phenotype similarity. Although the method is not designed to uniformly outperform all alternatives in every setting, its primary strength lies in adaptively combining complementary signals to improve phenotype prioritization across heterogeneous rare-variant scenarios.

## Materials and methods

### UK Biobank 500K dataset and genotype data quality control

We conducted our analysis using the UK Biobank 500K dataset [32, 33]. To reduce potential confounding due to population stratification, we restricted the study population to individuals of European ancestry. We applied multiple filtering criteria to ensure data quality and completeness, excluding individuals without hospital inpatient records, covariates information or whole-exome sequencing (WES) data. To mitigate bias arising from sample relatedness, we further removed related individuals using kinship coefficients provided by UK Biobank. Specifically, participants with first- or second-degree relationships were identified, and one individual was randomly retained from each related family. After applying these filters, the final study cohort consisted of 328,737 unrelated individuals of European ancestry. This curated dataset provides a homogeneous population with comprehensive phenotypic and genetic information, improving the robustness and interpretability of downstream analyses.

To ensure the quality of rare variant analyses, we performed stringent quality control (QC) on the WES data using PLINK [34]. Variants with a minor allele frequency (MAF) greater than 0.01 were removed, retaining only rare variants for subsequent analyses. We further excluded variants that deviated significantly from Hardy–Weinberg equilibrium (HWE; *p <* 1 × 10^−6^), which may indicate genotyping errors or residual population structure. Samples with missing sex information were also excluded. Following QC, we identified rare predicted loss-of-function (pLoF) variants, including stop-gain, stop-loss, startloss, frameshift insertions, frameshift deletions, frameshift substitution and essential splicing exonic variants, enabling a focused analysis of variants with a high likelihood of functional impact.

### Embedding models for ICD-10 codes, phecodes and individuals

To embed the ICD-10 codes in the UK Biobank (UKBB) dataset, we applied the Word2Vec model [11] to learn distributed representations of diagnosis codes. For each individual, we extracted the longitudinal sequence of ICD-10 codes from the electronic health records (EHR), treating each ICD-10 code as a token and each individual’s code trajectory as a sentence. The original temporal ordering of codes was preserved to retain contextual and positional information within the clinical history. We then trained the Word2Vec model on the full EHR corpus to obtain static embeddings for each ICD-10 code, which were subsequently used in downstream analyses.

Using the learned ICD-10 code embeddings from each model, we constructed both individual-level phenotype embeddings and phecode embeddings. For individual embeddings, we computed the average of the embeddings corresponding to all ICD-10 codes observed in an individual’s EHR:

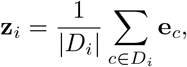

where *D*_*i*_ denotes the set of ICD-10 codes recorded for individual *i*, |*D*_*i*_| denotes the number of recorded ICD-10 codes for individual *i*, and **e**_*c*_ represents the embedding of ICD-10 code *c*.

For phecode embeddings, we aggregated the embeddings of ICD-10 codes mapped to each phecode using frequency-based weighting:

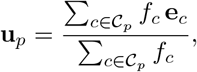

where 𝒞_*p*_ denotes the set of ICD-10 codes mapped to phecode *p*, and *f*_*c*_ is the frequency of code *c* in the EHR dataset. These procedures yield consistent individual-level and phecode-level embeddings for downstream phenome-wide analyses.

### EmbedPheScan

Within the EmbedPheScan framework, the objective is to perform a phenome-wide scan for a given gene in order to identify phenotypes most strongly associated with that gene. We focus on rare loss-of-function (LoF) variants, as such variants are more likely to have substantial phenotypic effects compared with common or missense variants when a gene is pathogenic. Accordingly, for each gene, we first identify individuals carrying rare LoF variants and aggregate their phenotypic information to characterize the gene’s phenotypic profile.

Specifically, let *C*_*g*_ denote the set of individuals carrying rare LoF variants in gene *g*. We construct a gene-level phenotype embedding by averaging the individual-level phenotype embeddings of these carriers:

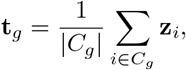

where **z**_*i*_ represents the phenotype embedding of individual *i*, |*C*_*g*_| denotes the number of individuals carrying rare LoF variants in gene *g*. This gene embedding summarizes the shared phenotypic characteristics of rare LoF variant carriers.

Next, we perform a phenome-wide scan by computing the similarity between the gene-level phenotype embedding and each phecode embedding. For each phecode *p* with embedding **u**_*p*_, we calculate the cosine similarity:

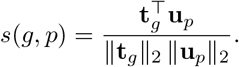

To assess the statistical significance of the similarity scores produced by EmbedPheScan, we adopt a sampling-based strategy to assess statistical significance. For each gene *g*, we generate an empirical reference distribution that preserves the carrier count while breaking any gene–phenotype relationship.

Specifically, let *n*_*g*_ = |*C*_*g*_| denote the number of rare LoF carriers for gene *g*. We repeatedly (10,000 times) sample without replacement a set of *n*_*g*_ individuals from the full cohort to form pseudo-carrier groups that mimic a null gene. For each random group *b* = 1,..., *B*, we compute a null gene embedding

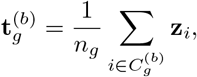

and evaluate its cosine similarity with every phecode embedding **u**_*p*_, yielding null similarity scores *s*^(*b*)^(*g, p*). This procedure produces an empirical null distribution for each gene–phecode pair that properly reflects the carrier count and the geometry of the embedding space.

We then standardize the observed similarity score *s*(*g, p*) using the empirical mean and standard deviation of the null distribution:

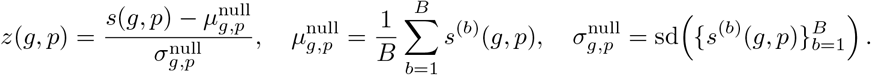

Under the null hypothesis of no gene–phenotype association, the standardized statistic is approximately standard normal. We therefore obtain a one-sided empirical p-value as

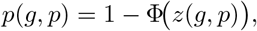

where Φ (*·*) denotes the cumulative distribution function of the standard normal distribution. These p-values are subsequently used to produce the final phenome-wide ranking and downstream multiple-testing analyses.

This resampling-based calibration ensures that significance estimates properly account for carrier sample size, phenotype prevalence structure, and embedding geometry, thereby providing well-calibrated inference for EmbedPheScan across genes with widely varying numbers of rare variant carriers.

### Conventional PheWAS

Besides the EmbedPheScan framework, we conducted a rare LoF carrier-based phenome-wide association study (PheWAS) for each gene. Let *g* denote a gene and let 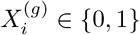 indicate whether individual *i* carries a rare LoF variant in gene *g*. For each phecode *p*, we define the binary phenotype indicator *Y*_*ip*_ ∈ {0, 1} based on the presence of the corresponding phecode in the individual’s EHR. We fit a covariate-adjusted logistic regression model

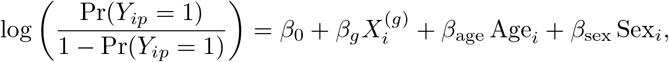

where *β*_*g*_ represents the log-odds ratio associated with LoF carrier status. The Wald test *p*-value for *β*_*g*_ is reported as the conventional PheWAS association statistic. When the logistic model failed to converge, we instead applied Fisher’s exact test on the corresponding contingency table.

Across all phecodes tested for a gene, we applied Benjamini–Hochberg false discovery rate (FDR) control to account for multiple comparisons. Phecodes were ranked by their conventional PheWAS *p*-values for downstream evaluation.

### EA-PheWAS

To leverage complementary strengths of embedding-based and regression-based association signals, we combine the p-values from EmbedPheScan and the conventional conventional PheWAS using the Aggregated Cauchy Association Test (ACAT) [35]. ACAT provides a powerful and computationally efficient framework for combining potentially dependent p-values while maintaining proper type I error control.

For a given gene–phecode pair (*g, p*), let *p*_embed_(*g, p*) denote the p-value obtained from EmbedPheScan, and let *p*_raw_(*g, p*) denote the corresponding p-value from the conventional PheWAS. ACAT transforms each p-value using the Cauchy quantile function and forms a weighted sum:

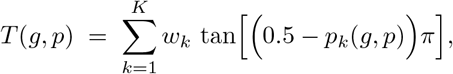

where *p*_*k*_(*g, p*) indexes the input p-values (here *K* = 2), and *w*_*k*_ ≥ 0 are nonnegative weights satisfying ∑_*k*_ *w*_*k*_ = 1. In our primary analysis, we use equal weights *w*_1_ = *w*_2_ = 0.5.

Under the global null hypothesis, the statistic *T* (*g, p*) approximately follows a standard Cauchy distribution regardless of the dependence structure among the input p-values. The combined ACAT p-value is therefore computed as

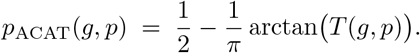

This aggregation strategy has several advantages in our setting. The embedding-based test in EmbedPheScan is sensitive to distributed phenotypic similarity patterns, while conventional PheWAS is optimized for single-phenotype regression signals. EA-PheWAS adaptively amplifies evidence when either method is informative and remains well calibrated when one component is weak or conservative. Consequently, the combined statistic improves robustness across genes with heterogeneous genetic architectures and phenotype manifestations.

The resulting *p*_ACAT_(*g, p*) values are used alongside the individual methods for ranking phenotypes and for downstream performance evaluation.

### Simulations to assess false-positive rates

We assessed type I error control through a gene-level simulation that preserves the empirical structure of the UK Biobank data.

For each simulation replicate, we constructed a null gene by randomly sampling a set of individuals and treating them as pseudo-carriers. The sample size was fixed to mimic realistic rare LoF carrier counts observed in UK Biobank. Specifically, we considered three carrier-count settings corresponding to the 25%, 50%, and 75% quantiles of the LoF carrier distribution, which are approximately 50, 120, and 300 carriers per gene, respectively. Within each setting, we generated multiple independent null replicates.

For every simulated null gene, we performed a full phenome-wide scan using three approaches: EmbedPheScan, conventional PheWAS, and EA-PheWAS. This produced a set of p-values across all phecodes for each replicate.

We evaluated calibration using two diagnostics. First, we generated quantile–quantile (QQ) plots comparing the observed p-value distribution to the expected uniform distribution under the null. Second, we computed the empirical type I error rate at a nominal significance level *α* as

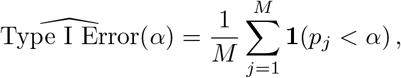

where *p*_*j*_ denotes the p-value from the *j*-th null test and *M* is the total number of tests across all replicates.

This framework allows direct assessment of whether each method maintains appropriate calibration across realistic rare-variant carrier regimes.

### Evaluation analysis

The evaluation analysis aims to assess the ability of EA-PheWAS to recover clinically relevant phenotypes for a given gene based on established reference information. In current PheWAS studies, the most widely used approach for prioritizing gene-associated phenotypes is regression-based association testing. Accordingly, we compared the performance of EA-PheWAS with both EmbedPheScan and the regression-based PheWAS framework (conventional PheWAS) as baselines.

For reference phenotype construction, we leveraged the Human Phenotype Ontology (HPO) database to obtain clinically validated phenotypes associated with each gene. For a given gene, we extracted the corresponding set of HPO terms curated in the HPO database. Because HPO terms represent well-defined and specific phenotypic abnormalities, they can be mapped to phecodes with minimal information loss [6]. This mapping yields a gene-specific list of clinically validated phecodes, which we treat as the reference phenotype set.

We evaluated model performance using hit precision, which measures the proportion of top-ranked phenotypes that appear in the reference set. Specifically, for a given gene *g*, let 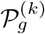 denote the set of top-*k* phecodes ranked by a method, and let 𝒢_*g*_ denote the reference phecode set derived from HPO. Hit precision at *k* is defined as

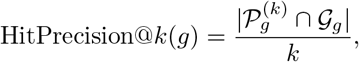

where *k* ∈ {5, 10, 15}. Higher hit precision indicates better recovery of clinically relevant phenotypes.

To summarize performance across genes, we compute the average hit precision by averaging HitPrecision@*k* over all evaluated genes. Let 𝒮 denote the set of genes included in the evaluation. The average hit precision is defined as

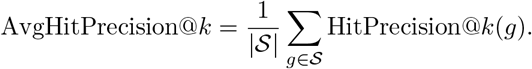

We also evaluated recall, which measures the proportion of ground truth phenotypes that are identified as significant by each method. For a given gene *g*, let 𝒮_*g*_ denote the set of phecodes identified as significant by a method after multiple testing correction, and let 𝒢_*g*_ denote the ground truth phecode set derived from HPO. The recall for gene *g* is defined as

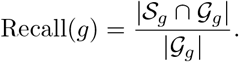

The average recall rate across genes is then calculated as

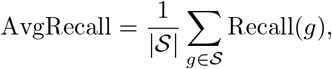

where 𝒮 denotes the set of genes included in the evaluation.

To further compare methods, we define the beat rate, which measures how often one method consistently achieves equal or higher hit precision than another method across multiple ranking thresholds. For two methods *A* and *B*, the beat rate is defined as

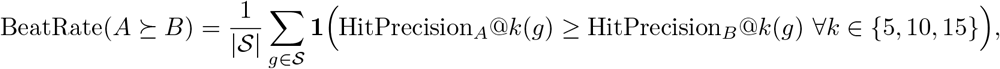

where **1**(*·*) denotes the indicator function.

Using this definition, we report the beat rate of EA-PheWAS compared with EmbedPheScan, the regression-based PheWAS baseline, and the proportion of genes for which EA-PheWAS simultaneously outperforms both methods under the same criterion.

## Code Availability

Code for EA-PheWAS is publicly available at https://github.com/JJJJJasonZheng/EA-PheWAS.

## Acknowledgments

This research was conducted using data from the UK Biobank Resource (Application ID: 29900). We gratefully acknowledge the UK Biobank participants, whose generous contributions made this study possible.

## Notes

### Competing Interest Statement

The authors have declared no competing interest.

