## Supplement_pdf for "EA-PheWAS: Integrating Phenotype Embeddings with PheWAS for Enhanced Gene-Phenotype Discovery"

### Supporting information

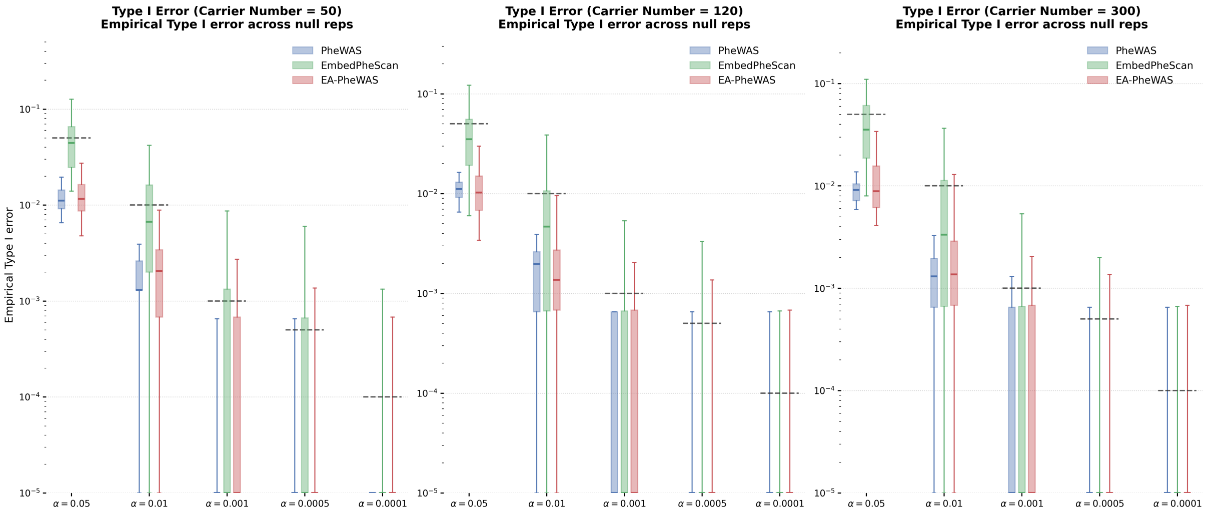


**Fig S1. Empirical type I error across null simulations.** Empirical type I error rates for conventional PheWAS, EmbedPheScan, and EA-PheWAS across simulation replicates are shown at multiple nominal significance thresholds for carrier counts of 50, 120, and 300. Dashed lines denote the nominal error rates.

**Table S1. Average hit precision across genes.** Mean, median, standard deviation, and number of evaluated genes are shown for conventional PheWAS, EmbedPheScan, and EA-PheWAS at Top-5, Top-10, and Top-15 ranked phenotypes. Hit precision is defined as the proportion of top-$k$ prioritized phenotypes that are also present in the HPO reference set.

**Table S2. Recall improvement relative to conventional PheWAS.** Mean and median absolute recall differences are shown for EmbedPheScan and EA-PheWAS relative to conventional PheWAS across multiple FDR thresholds. Recall is defined as the proportion of HPO reference phenotypes recovered among the significant phenotypes identified for each gene.

**Table S3. Average hit precision across HI genes.** Mean, median, standard deviation, and number of evaluated genes are shown for conventional PheWAS, EmbedPheScan, and EA-PheWAS at Top-5, Top-10, and Top-15 ranked phenotypes. Hit precision is defined as the proportion of top-$k$ prioritized phenotypes that are also present in the HPO reference set.

**Table S4. Average hit precision across tissue specific genes.** Mean, median, standard deviation, and number of evaluated genes are shown for conventional PheWAS, EmbedPheScan, and EA-PheWAS at Top-5, Top-10, and Top-15 ranked phenotypes. Hit precision is defined as the proportion of top-$k$ prioritized phenotypes that are also present in the HPO reference set.

**Table S5. Average hit precision across non-tissue genes.** Mean, median, standard deviation, and number of evaluated genes are shown for conventional PheWAS, EmbedPheScan, and EA-PheWAS at Top-5, Top-10, and Top-15 ranked phenotypes. Hit precision is defined as the proportion of top-$k$ prioritized phenotypes that are also present in the HPO reference set.

**Table S6. Phenome-wide association results for *PKD1* under EA-PheWAS.** The table lists phecodes prioritized for *PKD1* together with association statistics from conventional PheWAS, embedding-based similarity statistics from EmbedPheScan, and aggregated ACAT p-values from EA-PheWAS. Specifically, *phecode* denotes the tested phenotype code; *p_lr* denotes the conventional PheWAS p-value from regression-based association testing; *phecode_des* gives the phenotype description; *beta_icd* and *se_icd* denote the estimated effect size and standard error from conventional PheWAS; *n_case* denotes the total number of phenotype cases; *p_embed* denotes the embedding-based p-value from EmbedPheScan; *centered_similarity* denotes the similarity score centered relative to the empirical null distribution; and *p_acat* denotes the aggregated p-value from EA-PheWAS. Results are ordered by *p_acat*.

**Table S7. Phenome-wide association results for *PKD2* under EA-PheWAS.** The table lists phecodes prioritized for *PKD2* together with association statistics from conventional PheWAS, embedding-based similarity statistics from EmbedPheScan, and aggregated ACAT p-values from EA-PheWAS. Specifically, *phecode* denotes the tested phenotype code; *p_lr* denotes the conventional PheWAS p-value from regression-based association testing; *phecode_des* gives the phenotype description; *beta_icd* and *se_icd* denote the estimated effect size and standard error from conventional PheWAS; *n_case* denotes the total number of phenotype cases; *p_embed* denotes the embedding-based p-value from EmbedPheScan; *centered_similarity* denotes the similarity score centered relative to the empirical null distribution; and *p_acat* denotes the aggregated p-value from EA-PheWAS. Results are ordered by *p_acat*.

**Table S8. Phenome-wide association results for *NF1* under EA-PheWAS.** The table lists phecodes prioritized for *NF1* together with association statistics from conventional PheWAS, embedding-based similarity statistics from EmbedPheScan, and aggregated ACAT p-values from EA-PheWAS. Specifically, *phecode* denotes the tested phenotype code; *p_lr* denotes the conventional PheWAS p-value from regression-based association testing; *phecode_des* gives the phenotype description; *beta_icd* and *se_icd* denote the estimated effect size and standard error from conventional PheWAS; *n_case* denotes the total number of phenotype cases; *p_embed* denotes the embedding-based p-value from EmbedPheScan; *centered_similarity* denotes the similarity score centered relative to the empirical null distribution; and *p_acat* denotes the aggregated p-value from EA-PheWAS. Results are ordered by *p_acat*.

**Table S9. Phenome-wide association results for *FBN1* under EA-PheWAS.** The table lists phecodes prioritized for *FBN1* together with association statistics from conventional PheWAS, embedding-based similarity statistics from EmbedPheScan, and aggregated ACAT p-values from EA-PheWAS. Specifically, *phecode* denotes the tested phenotype code; *p_lr* denotes the conventional PheWAS p-value from regression-based association testing; *phecode_des* gives the phenotype description; *beta_icd* and *se_icd* denote the estimated effect size and standard error from conventional PheWAS; *n_case* denotes the total number of phenotype cases; *p_embed* denotes the embedding-based p-value from EmbedPheScan; *centered_similarity* denotes the similarity score centered relative to the empirical null distribution; and *p_acat* denotes the aggregated p-value from EA-PheWAS. Results are ordered by *p_acat*.
